# The complete connectome of a learning and memory center in an insect brain

**DOI:** 10.1101/141762

**Authors:** Katharina Eichler, Feng Li, Ashok Litwin-Kumar, Youngser Park, Ingrid Andrade, Casey M. Schneider-Mizell, Timo Saumweber, Annina Huser, Claire Eschbach, Bertram Gerber, Richard D. Fetter, James W. Truman, Carey E. Priebe, L. F. Abbott, Andreas S. Thum, Marta Zlatic, Albert Cardona

**Affiliations:** HHMI Janelia Research Campus, 19700 Helix Dr., Ashburn, VA 20147; Department of Biology, University of Konstanz, 78464 Konstanz, Germany; Department of Neuroscience, Columbia University; Department of Applied Mathematics and Statistics, Whiting School of Engineering, Johns Hopkins University; Abteilung Genetik von Lernen und Gedächtnis, Leibniz Institut für Neurobiologie (LIN), 39118 Magdeburg, Germany; Otto von Guericke Universität Magdeburg, Institut für Biologie, Verhaltensgenetik, Universitätsplatz 2, 39106 Magdeburg, Germany; Center for Behavioral Brain Sciences (CBBS), 39106 Magdeburg, Germany; Department of Physiology and Cellular Biophysics, Columbia University; Department of Zoology, University of Cambridge; Department of Physiology, Development and Neuroscience, University of Cambridge

## Abstract

Associating stimuli with positive or negative reinforcement is essential for survival, but a complete wiring diagram of a higherorder circuit supporting associative memory has not been previously available. We reconstructed one such circuit at synaptic resolution, the *Drosophila* larval mushroom body, and found that most Kenyon cells integrate random combinations of inputs but a subset receives stereotyped inputs from single projection neurons. This organization maximizes performance of a model output neuron on a stimulus discrimination task. We also report a novel canonical circuit in each mushroom body compartment with previously unidentified connections: reciprocal Kenyon cell to modulatory neuron connections, modulatory neuron to output neuron connections, and a surprisingly high number of recurrent connections between Kenyon cells. Stereotyped connections between output neurons could enhance the selection of learned responses. The complete circuit map of the mushroom body should guide future functional studies of this learning and memory center.

---

Massively parallel, higher-order neuronal circuits such as the cerebellum and insect mushroom body serve to form and retain associations between stimuli and reinforcement in vertebrates and higher invertebrates^1–6^. Although these systems provide a biological substrate for adaptive behavior, no complete synapse-resolution wiring diagram of their connectivity has been available to guide analysis and inspire understanding. The mushroom body (MB) is a higher-order parallel fiber system in many invertebrate brains, including hemimetabolous as well as holometabolous insects and their larval stages^6^. MB function is essential for associative learning in adult insects^1;3–5^ and in *Drosophila* larvae^1;7;8^, from the earliest larval stages onward^9^. Indeed, the basic organization of the adult and the larval MB and their afferent circuits is very similar; however, larvae have about an order of magnitude fewer neurons^7^. Thus, to systematically investigate the organizational logic of the MB, we used serial section electron microscopy (EM) to map with synaptic resolution the complete MB connectome in a first instar *Drosophila* larva (L1; Fig. 1a). L1 are foraging animals capable of all behaviors previously described in later larval stages^10^, including adaptive behaviors dependent on associative learning^7;9^ (Fig. 1b). Their smaller neurons enable fast EM imaging of the entire nervous system and reconstruction of complete circuits^11;12^.

**Figure 1.**
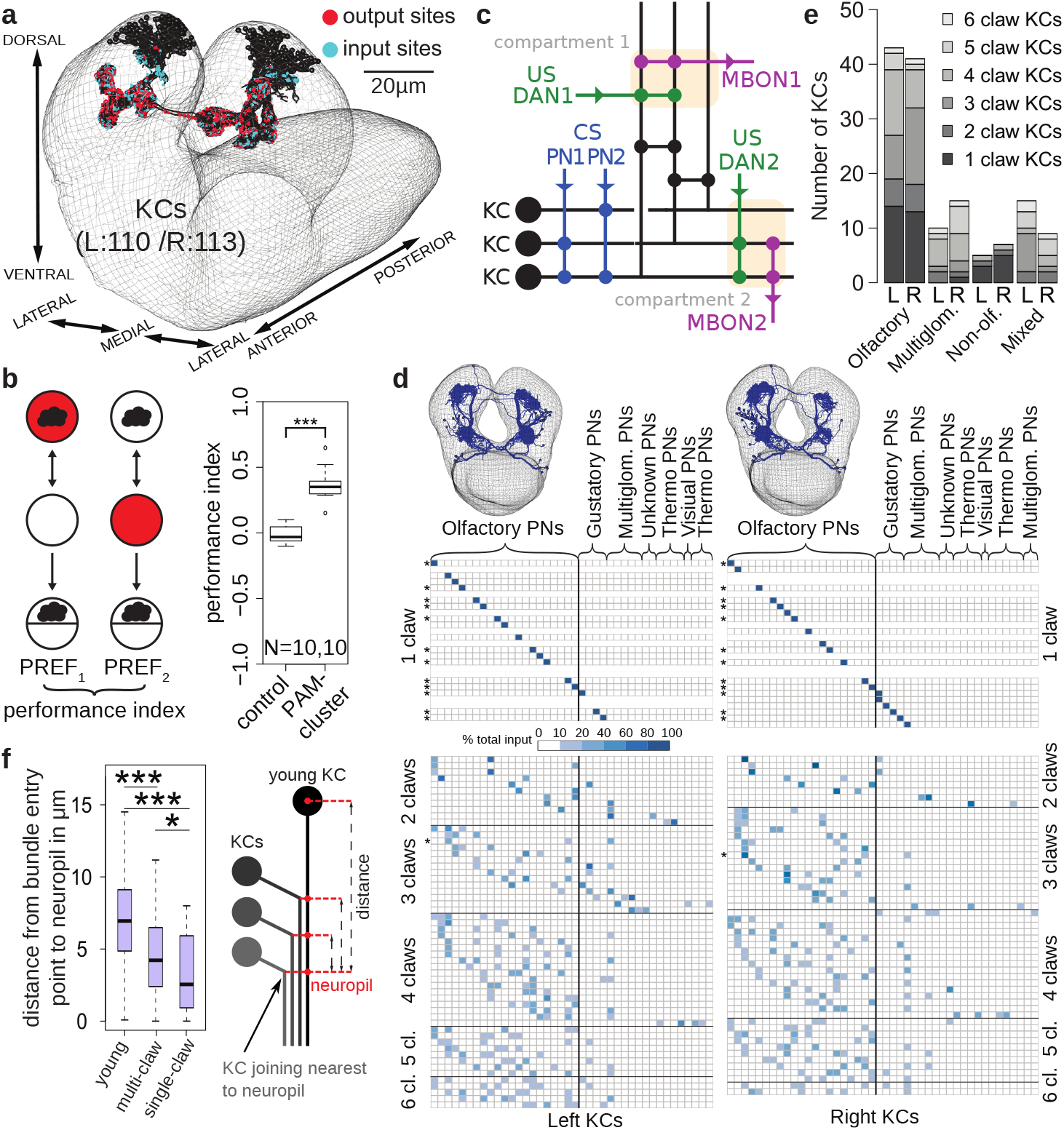
Mushroom bodies of a first instar *Drosophila* larva. **a** Kenyon cells (KCs) from a whole-CNS EM volume. **b** Associative learning in first instar larvae: in a Petri dish, we presented an odor (cloud) and red light, either paired (left) or unpaired (right), and computed the performance index. Control larvae (attP2; UAS-CsChrimson) receiving paired stimuli did not learn, whereas larvae in which optogenetic activation of dopaminergic (PAM cluster) neurons (GMR58E02-GAL4; UAS-CsCrimson) was paired with odor showed robust appetitive learning (*p* < 0.0001). c Diagram of the literature’s MB circuitry model. Projection neurons (PN) relay sensory stimuli to KC dendrites. MBON dendrites and MBIN (DAN, OAN and other) axons tile the parallel KC axons, defining compartments (*colored boxes*). MBINs signal reward or punishment, and KCs synapse onto output neurons (MBONs). **d** PN-to-KC connectivity, color-coded by % of inputs on KC dendrites. Uniglomerular olfactory PNs (Olfactory PNs) and other PNs synapse onto single-claw or multi-claw KCs. Stars indicate KCs with identical input patterns on the left and right hemispheres. For the PNs on the right of the black vertical lines, first 18 columns are left-right homologous PNs, last column (left) and last five columns (right) are hemisphere-specific PNs. **e** Number of KCs integrating inputs from uniglomerular olfactory, multiglomerular olfactory, non-olfactory or a mixture of these PN types. **f** Earlier-born KCs join the lineage bundle closer to the neuropil surface than later-born ones, meaning older KCs present less claws than younger ones. Distances span from the point where the KC joins the bundle to the joining point of the KC nearest to the neuropil. Differences between all groups are significant (*** p-values smaller than 0.0001; single-claw and multi-claw KC comparison p-value: 0.0237).

Models of sensory processing in many neural circuits feature neurons that fire in response to combinations of sensory inputs, generating a high-dimensional representation of the sensory environment^13^. The intrinsic MB neurons, the Kenyon cells (KCs), integrate in their dendrites inputs from combinations of projection neurons (PNs) that encode various stimuli, predominantly olfactory in both adult^1;4–6^, and larva^12;14^, but also thermal, gustatory and visual in adult^5;6;15^ and larva (reported here for the first time). Previous analyses in adults^16;17^ and larvae^14^ suggest that the connectivity between olfactory PNs and KCs is random, but they do not eliminate the possibility of some degree of bilateral symmetry, which requires access to the full PN → KC wiring diagram in both hemispheres.

The MB contains circuitry capable of associating reward or punishment with the representation of the sensory environment formed by KCs. KCs have long parallel axons that first run together forming the peduncle and then bifurcate, forming the so-called lobes, in both larvae^7^ and adults^1;4;6;18^. KCs receive localized inputs along their axonal fibers from dopaminergic as well as octopaminergic modulatory neurons (DANs and OANs, respectively) that define separate compartments. DANs and OANs have been shown to convey reinforcement signals in adult insects^3–6;19–24^ and larval *Drosophila*^7;8^. The dendrites of the mushroom body output neurons (MBONs) respect the DAN compartments in adult^5;18;25;26^ and larvae^9^. It has been shown in adult *Drosophila* that co-activation of KCs and DANs can associatively modulate the KC-MBON synapse^5;19;20;22;24;26–28^. Thus, the compartments represent anatomical and functional MB units where sensory input (KCs) is integrated with internal reinforcement signals (DANs/OANs) to modulate instructive output for behavioral control (MBONs). However, the synaptic connectivity of KCs, DAN/OANs, and MBONs at this crucial point of integration was previously unknown.

Furthermore, studies in adult *Drosophila* have shown that despite the compartmental organization of the MB, many MBONs interact with MBONs from other compartments, suggesting that the MBON network functions combinatorially during memory formation and retrieval^18;25;28^. However, a comprehensive account of all MB neuron connections is lacking. Thus, to provide a basis for understanding how the MB, a prototypical parallel fiber learning and memory circuit, functions as an integrated whole, we provide a full, synapse-resolution connectome of all MB neurons of an L1 *Drosophila* larva.

## Results

We reconstructed all the KCs in both brain hemispheres of an L1 *Drosophila* larva and identified all of their pre- and postsynaptic partners (Fig. 1a, d, Fig. 3a; Supp. Table 1). We found 223 KCs (110 on the left, 113 on the right), of which 145 are mature (73 on the left, 72 on the right). Immature KCs either lack or have tiny dendrites, and their axons terminate early with filopodia typical of axon growth cones. Every mature KC presents a well-developed dendrite, and its axon innervates all of the MB compartments. Although the number of KCs is different between the two sides (Fig. 1a), we found exactly 24 MBONs, 7 DANs, 2 paired and 2 unpaired OANs and 5 additional modulatory input neurons (which we call MBINs or mushroom body input neurons, a term we also use to refer to all the modulatory neurons collectively) of unknown neurotransmitter identity (Supp. Fig. 1a, Supp. Table 3). An additional GABAergic neuron, homologous to the APL neurons in the adult fly^29;30^, synapses reciprocally with all mature ipsilateral KCs (Ext. Data Fig. 1).

## PN inputs to the KCs

We identified all the sensory PNs and their connections onto KCs. KC dendrites have claw-like structures that wrap around PN axon boutons in the MB calyx^6^ (Ext. Data Fig. 2b). We identified input from 21 uniglomerular olfactory PNs on each side^12^, 5 and 7 multiglomerular PNs ^12^, and 14 and 16 non-olfactory PNs on the left and right sides, respectively. Non-olfactory PNs include thermal, visual, gustatory, mixed olfactory, and possibly other modalities (Fig. 1d, e; Supp. Table 1). A subset of mature KCs receives input from only one PN (single-claw KCs), while the remaining KCs (multi-claw KCs, as in the adult fly^17^) receive input from 2–6 PNs^14^ (Fig. 1d, Ext. Data Fig. 2c). Interestingly, single-claw KCs are born earlier than multi-claw KCs (Fig. 1f). KCs with different numbers of claws receive roughly the same number of synapses summed across claws (Ext. Data Fig. 2a). This suggests that multi-claw KCs may require input from multiple PNs to respond, assuring combinatorial selectivity^31;32^.

Two features of the PN → KC connectivity are immediately visible (Fig. 1d): the contrast between the ordered and apparently disordered connections onto single-claw KCs and multi-claw KCs, respectively, and the existence of KCs that do not receive any uni-glomerular PN input. Most KCs (77%) receive exclusively olfactory input from uni-or multi-glomerular PNs, while the others receive non-olfactory input (Fig. 1d, e). Prominent among these are two KCs per side that exclusively receive thermal information.

No structure is apparent in the olfactory PN input to multi-claw KCs (Fig. 1d; see Ext. Data Fig. 3b–e for further analysis), consistent with previous analyses in adults^16;17^ and larvae^14^. Structure was found, however, when all the PNs were included in the analysis, reflecting the presence of specialized non-olfactory KCs (Ext. Data Fig. 3a). We also determined that PN connections onto multi-claw KCs in the left and right hemispheres are statistically independent. Only one bilateral pair of multi-claw KCs receives input from the same set of homologous PNs (marked by asterisks in Fig. 1d), no more than predicted by a random model (*p* = 0.52; Ext. Data Fig. 3e). Such asymmetry has not been observed previously in the L1 where strongly connected presynaptic partners of identified neurons have always been seen to have left and right homologs^11;12^.

In contrast to the randomness of multi-claw KCs, the number of single-claw KCs is significantly greater than predicted by random models (Fig. 2a) and Fig. 1d shows clear structure in their wiring. If single-claw KCs sampled random PNs, we would expect to find about 5 pairs of single-claw KCs innervated by the same PN per hemisphere (“duplicates”). The data indicate zero left-hemisphere and one right-hemisphere duplicate (*p* < 0.002 andp < 0.005 in the random model, respectively). In contrast, the random model predicts that duplicates should be rare for multiclaw KCs, and the data reveal only two (*p* = 0.93). The fact that single-claw KCs appear earliest in development suggests that a top priority, initially, is to assure that a complete set of signals is relayed to the MBONs, which is not guaranteed with random wiring. Indeed, we calculated that 27 (left) and 44 (right) randomly wired multi-claw KCs, on average, were needed to achieve the same level of coverage of PN inputs as the 17 (left) and 19 (right) single-claw KCs (Methods).

**Figure 2.**
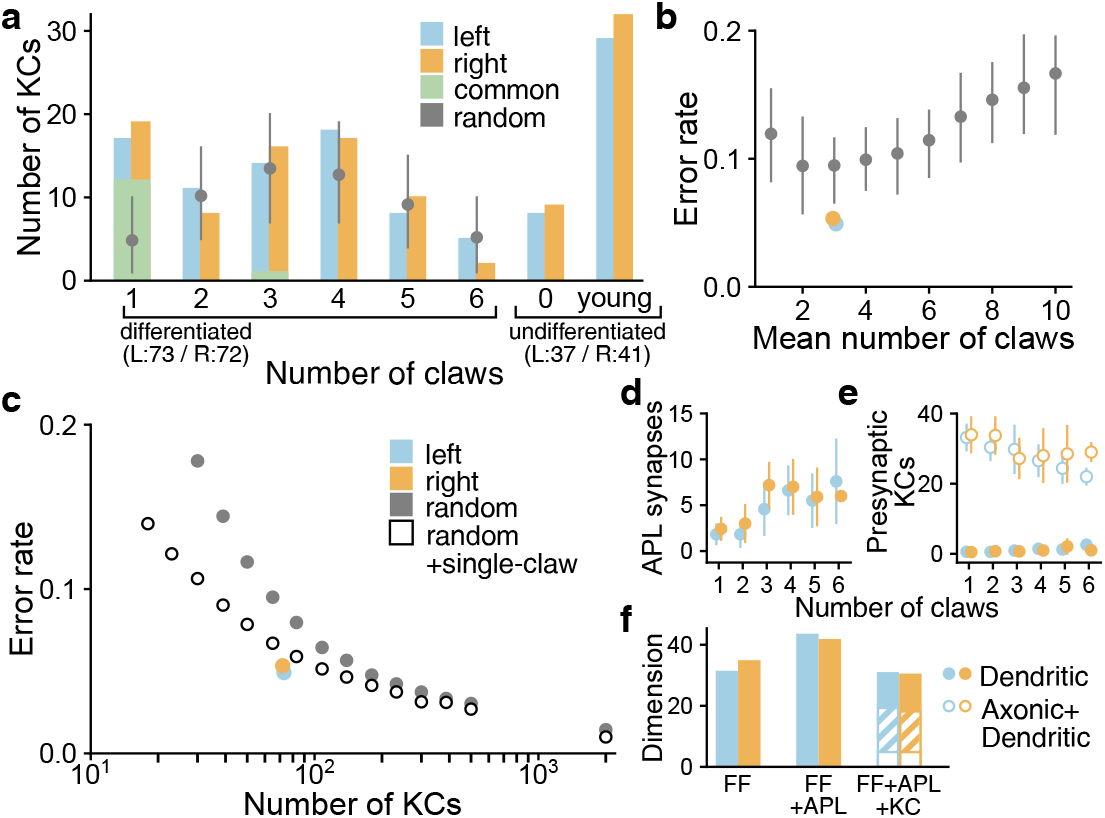
KC connectivity reduces redundancy and optimizes stimulus discrimination. **a** Distribution of KC claw numbers compared to random models. Random models have significantly fewer single-claw KCs (*p* < 10^−5^). Grey circles and lines denote mean and 95% confidence intervals. **b** Classification error rate of a readout of the KC representation trained on a stimulus discrimination task. Observed connectivity (blue and orange) is compared to random models (grey) in which KCs have different distributions of average claw numbers. **c** Average performance of models with purely random connectivity (grey) or random multi-claw plus non-random single-claw KCs (white). S.E.M. is smaller than the marks. **d** Number of APL-to-KC synapses, which is correlated with claw number. **e** Number of presynaptic KC-to-KC connections, which is inversely related to postsynaptic claw number. Dendro-dendritic (filled circles) or all synapses (open circles) are shown. **f** Dimension of KC representation in models with only feedforward PN-to-KC connections (FF), with APL-mediated inhibition (FF+APL), or with inhibition and excitatory KC-to-KC connections (FF+APL+KC). Dimension is slightly reduced by dendritic KC-to-KC connections (right, filled bars) but strongly reduced by axonic KC-to-KC connections (open bars). Facilitatory axonic KC-to-KC connections (hatched bars) yield an intermediate reduction.

## PN → KC connectivity optimizes the KC odor representation for associative learning

The lack of duplication in PN → KC connections suggests that MB wiring is well-suited to promote KC diversity. We hypothesized that this diversity produces a high-dimensional odor representation that supports stimulus discrimination. To test this idea, we developed a model in which KCs produce sparse responses to random combinations of odor-evoked PN activity (Methods). We compared the performance on a stimulus classification task of a model MBON in networks with the observed PN → KC connectivity and completely randomly connected models with varying degrees of connectivity (see Methods). Fully random networks have few of the single-claw KCs observed in the reconstruction (Fig. 2a).

The observed connectivity leads to performance superior to the average performance of purely random model networks (Fig. 2b). Motivated by the observation that unique single-claw KCs are born early in MB development (Fig. 1f), we hypothesized that their presence may be particularly important when the number of KCs is small. We therefore compared the performance of networks with only randomly connected KCs to that of networks with the same total number of KCs but containing a subpopulation of unique single-claw KCs. In networks with few KCs, the presence of single-claw KCs substantially improves performance (Fig. 2c). As additional KCs are added, the advantage of single-claw KCs diminishes. For networks constructed using estimates of adult *Drosophila* KC connectivity, where single-claw KCs have not been identified^17^, unique single-claw KCs provide only a small benefit.

## Inhibitory KC interactions via the APL neuron

The MB is innervated by the GABAergic APL neuron in the adult fly^29^ and in the larva^30^. APL synapses reciprocally with all mature KCs (Ext. Data Fig. 1a). Although APL receives most of its input from multi-claw KCs, individual single-claw KCs typically have more synapses onto APL dendrites than multi-claw KCs (Ext. Data Fig. 1b,c). Conversely, APL connects more strongly to KCs with more claws (Fig. 2d, Ext. Data Fig. 1c). We extended our model to include APL-mediated feedback inhibition and assessed its effect on the dimension of the KC representation, which quantifies the level of decorrelation of KC responses^33^ (see Methods). The addition of recurrent APL inhibition increases the dimension by approximately 30% (Fig. 2f), consistent with the proposed role of APL in maintaining sparse, decorrelated KC responses^30;34;35^. The increased inhibition received by multi-compared to single-claw KCs may reflect a greater need for decorrelation of their responses given the larger overlap of their PN inputs.

## Recurrent KC interactions

We next examined whether KCs directly interact with each other, a possibility suggested by previous studies in *Drosophila* and other species ^6^. We found that, on average, 60% of the synapses received by KCs come from other KCs and 45% of KC output synapses are onto other KCs (Fig. 2e, Ext. Data Fig. 2d–g). Most KC → KC synapses are axo-axonic and located in the peduncle and MB lobes; a much smaller fraction are dendro-dendritic and located in the calyx (Ext. Data Fig. 2h). The largest number of KC → KC connections occurs between the two thermosensory KCs (Supp. Table 1), but these KCs also make large numbers of connections to olfactory and visual KCs. Single-claw KCs make more recurrent connections, on average, than multi-claw KCs, and both single- and multi-claw KCs tend to connect reciprocally to KCs of the same type (Ext. Data Fig. 4a–d). However, we found no relationship between the similarity of the PN inputs to KC pairs and the number of KC → KC synapses formed between them (correlation coefficients 0.006 and −0.02 for left and right hemispheres; *p* > 0.3, comparison to shuffled network).

KCs have been shown to be cholinergic in the adult^36^, so it is likely that KC → KC connections are depolarizing. Even when strong dendro-dendritic KC connections are added to our model, dimension only decreases slightly (Fig. 2f). Additional subthreshold facilitatory axo-axonic KC → KC connections also have a weak effect (Fig. 2f, hatched bars). However, if the axo-axonic connections we model are stronger than this, the dimension collapses due to the large correlations introduced by this recurrence (Fig. 2f, open bars). While our model does not reveal a functional role for KC → KC connections, it is possible that processes that we did not model, such as experience-dependent modulation of their synaptic strengths, may modify the representation to favor either behavioral discrimination or generalization^37^. Further characterization of these connections is needed to assess this hypothesis.

## MBINs and MBONs

We next identified the MBONs and modulatory neurons (MBINs) associated with every MB compartment. Each compartment is innervated by 1–5 MBONs and 1–3 MBINs (most often DANs and OANs), except for the shaft compartment of the medial lobe that lacks MBINs in L1 larvae (but has them in L3;^8^). Most MBONs and MBINs innervate a single MB compartment (although most MBINs present a bilateral axon, innervating both the left and right homologous compartments), with few exceptions (MBIN-l1 and 5/24 MBONs; Fig. 3d, Ext. Data Fig. 5). Most MBINs (over 80%) are primarily presynaptic to other MB neurons, but some allocate 50% or more of their outputs to non-MB neurons (Ext. Data Fig. 6a, Supp. Table 2b). We also found that larger MBIN axons make more synapses onto KCs, and likewise for KC synapses onto larger MBON dendrites (Ext. Data Fig. 6b, c).

**Figure 3.**
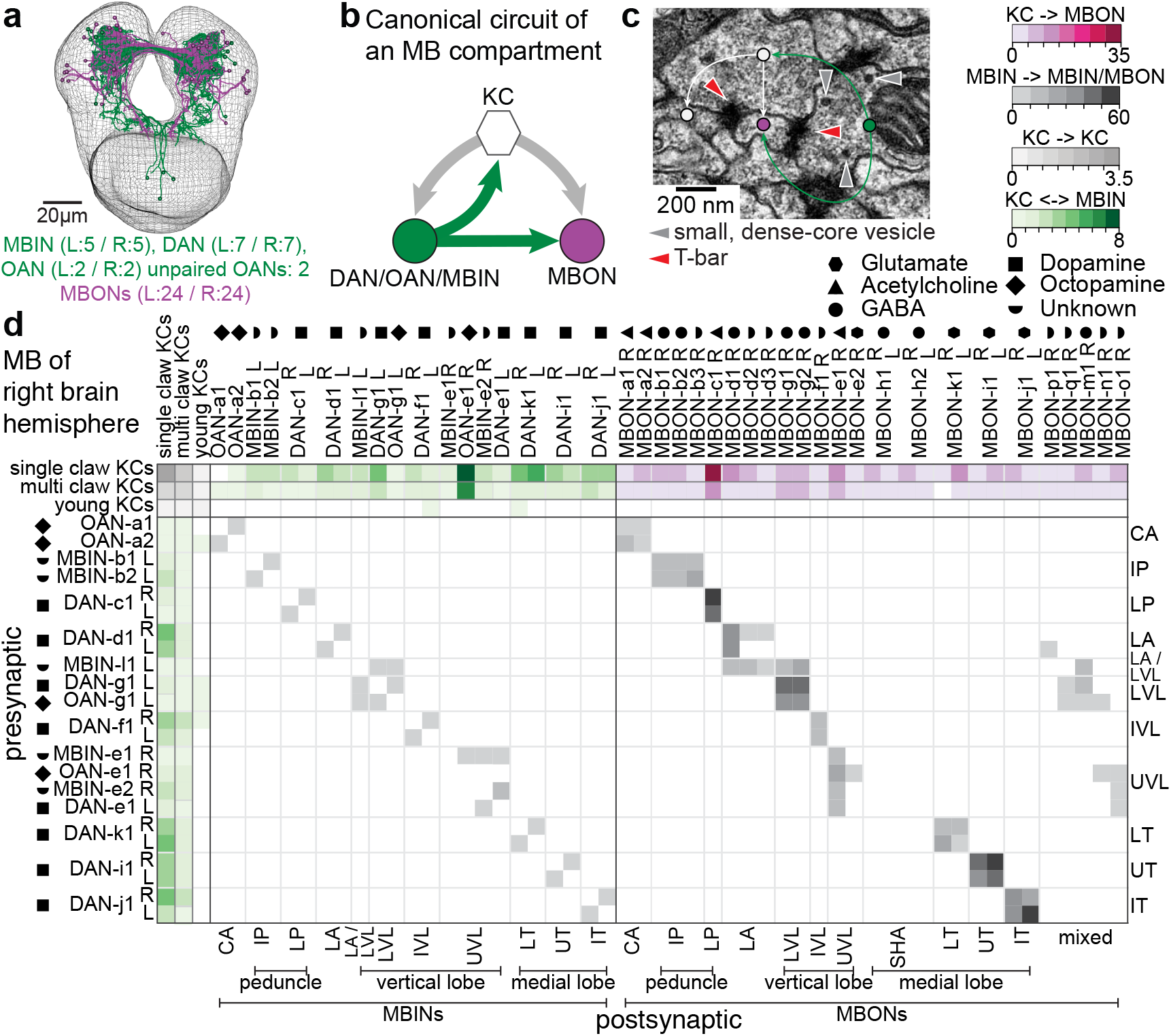
A canonical circuit in every mushroom body compartment. **a** EM-reconstructed MBIN (DANs, OANs and others) and MBONs. **b** Canonical circuit present in every MB compartment, with previously unknown KCs synapses onto MBINs, and from these onto MBONs. **c** Example of an MBIN (*green dot*) synapse with a KC (*white dot*) and an MBON (*magenta dot*). The same KC is also presynaptic to the MBON in close proximity. Dense- and clear-core vesicles are present near the DAN presynaptic site. **d** The connectivity matrix between KCs, MBINs, and MBONs of the right-hemisphere MB shows specific, compartment-centric synapses among cells types. Each entry represents the number of synapses from a row to a column (values are averaged for KCs). Note absence of DAN in SHA (develops later in larval life). MBINs synapse only onto MBONs innervating their compartment, and axo-axonically onto same-compartment MBINs. Note multi-compartment MBONs in the vertical lobe and LA.

Antibody labeling of GFP-labeled neurons showed that seven MBINs are dopaminergic (DAN-c1, d1, f1, g1, i1, j1 and k1; Ext. Data Fig. 5, four are octopaminergic (OAN-a1, -a2, -e1 and -g1), three others are neither dopaminergic nor octopaminergic (MBIN-e1, -e2 and -l1), and two were not technically accessible (MBIN-b1 and -b2) (Supp. Fig. 2a). MBONs can be glutamatergic, cholinergic or GABAergic (Supp. Fig. 2b), the same set of neurotransmitters seen for adult MBONs^18^ (Supp. Table 3).

## A canonical circuit motif in each MB compartment

EM reconstruction of MBONs and MBINs revealed a canonical circuit motif (Fig. 3b, d and Ext. Data Fig. 5) that appears in every compartment, independent of MBIN or MBON neurotransmitter (except in the shaft compartment that lacks modulatory input in L1). In this motif, KCs synapse onto MBONs, as previously shown in adult *Drosophila* ^5;6;24;36^ and bees and locusts^3;6;38^. However, we identified two unexpected connection types. The first are numerous KC → MBIN connections (Fig. 3d, Ext. Data Fig. 5). Second, a sizable fraction of MBIN presynaptic sites (which are polyadic) each simultaneously contacts both KCs and MBONs, with generally at least one of the postsynaptic KCs synapsing onto one of the postsynaptic MBONs within less than a micron of the MBIN-KC synapse (Fig. 3c). Thus, MBINs synapse both onto the pre- and postsynaptic side of many KC → MBON synapses. For comparison, MBONs receive on average 3.44% of their input from individual MBINs and 1.3% from an individual KC. If only 5% of the 73 mature KCs are active in response to a given odor, as has been shown in the adult^31^, then active KCs correspond to on average ca. 4.8% of the total synaptic inputs to MBONs, very similar to the % of MBIN input.

MBINs convey reinforcement signals and are thought to modulate the efficacy of KC → MBON connections through volume-release in the vicinity of KC presynaptic terminals^1;3–6;24–27^ and as expected, we observed MBIN axon boutons containing dense-core vesicles (Ext. Data Fig. 7). We also observed dense-core vesicles in addition to clear-core vesicles in 1/3 of KCs (Ext. Data Fig. 7), consistent with findings in the adult that many KCs corelease sNPF peptide with acetylcholine^36^.

Our EM reconstruction also revealed MBIN synapses containing clear vesicles (Ext. Data Fig. 7 and Fig. 3c) indistinguishable from those that release classical neurotransmitters^11^. All MBIN terminals with clear-core vesicles contained dense-core vesicles, mostly within presynaptic boutons (Ext. Data Fig. 8). This suggests that MBINs may have two concurrent modes of action onto KCs and MBONs: activation via synaptic release (of dopamine, octopamine, or fast neurotransmitters;^39^) and neuromodulation via volume release. These two modes, coupled with the diverse connection types among KCs, MBINs, and MBONs, may provide a powerful and flexible substrate for associative learning.

## Heterogeneous KC → MBON/MBIN connections in multiple compartments

Previous studies in adult *Drosophila* have shown that different lobes and compartments within the lobes are involved in forming different types of memories^1;4;5;19–22;25;26;28;40;41^. Functional studies in larvae^8;42^ also suggest that vertical and medial lobes may be implicated in distinct types of memory formation (aversive and appetitive, respectively). Our EM study shows that all mature KCs make synaptic connections with MBONs and MBINs (DANs, OANs, and others) in both the vertical and medial MB lobes (Ext. Data Fig. 9a-c, Ext. Data Fig. 10a-c and Supp. Table 4a, b). Furthermore, individual MBONs are innervated by between 40% to almost all of the KCs, with an average of 70% (Ext. Data Fig. 9a). This extensive innervation provides MBONs access to the high-dimensional KC representation and suggests that individual KCs may be involved in the formation and storage of associations involving multiple stimuli and valences, as suggested for the adult^1;4;5^ (but see^43^).

Our EM reconstruction reveals that in the larva, the axon terminals of all MBINs overlap with all KCs within a compartment and could potentially connect to all KCs, unlike in the adult^18;43^. Nevertheless, only subsets of KCs synapse onto either the MBIN or the MBON, or both, within a given compartment, with a broad distribution in the number of synaptic contacts (Ext. Data Fig. 9d, e, Ext. Data Fig. 10d, e and Supp. Table 4a, b). Estimating connection strength using synapse number, distinct subsets of KCs synapse strongly with MBINs and MBONs in distinct compartments (Ext. Data Fig. 9c, Ext. Data Fig. 10c, Supp. Table 4a, b and Supp. Table 1). This could arise from an innately broad distribution of synapse strengths or activity-dependent changes in synapse number. Either way, this heterogeneity implies that distinct MBONs and MBINs respond to heterogeneous combinations of active KCs.

## Comprehensive profile of MBON inputs

EM reconstruction and synaptic counting provides a comprehensive view of the signals carried by the MBONs (Fig. 4a, b and Supp. Table 2a). In general, MBONs are among the neurons in the L1 larval brain receiving the largest number of inputs, with a median of 497 and a maximum of about 1500 postsynaptic sites. The median for other neurons reconstructed so far is around 250 in L1^11;12^.

**Figure 4.**
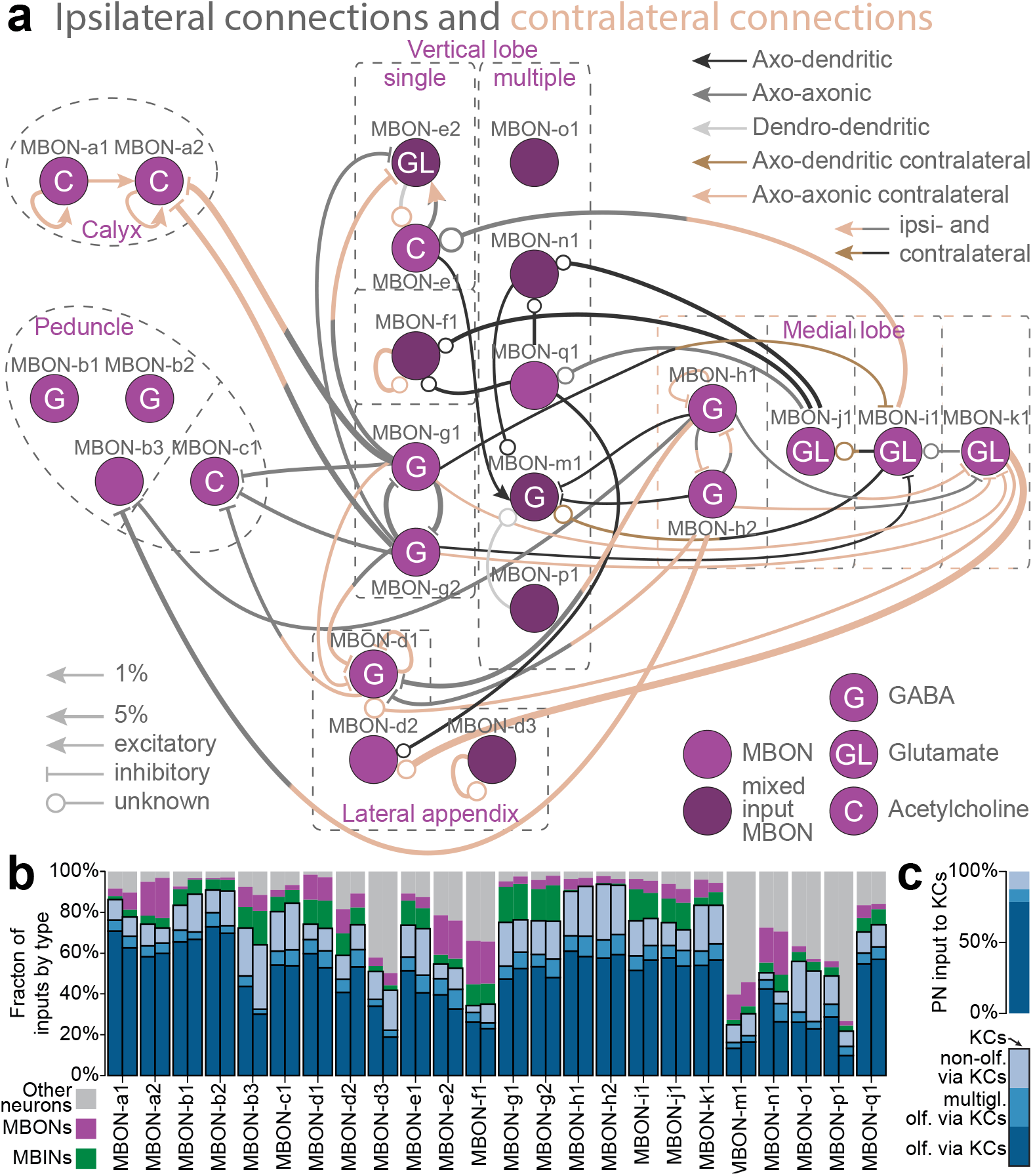
MBON inputs and circuits. **a** In the diagram, MBONs are in the compartments innervated by their dendrites. Most connections among MBONs are axo-axonic, and fewer are axo-dendritic. Few MBONs avoid synapsing to others. Most inter-lobe connections are mediated by GABAergic (MBON-g1, g2, h1, h2) and glutamatergic (MBON-i1, j1, k1) MBONs, potentially providing a substrate for lateral inhibition between compartments of opposite valence. **b** Fraction of MBON inputs by neuron type. Left and right homologous MBONs are shown adjacent. Only some vertical lobe and LA MBONs get less than 80% of their input from MB neurons and less than 60% from KCs. Almost all multi-compartment MBONs (MBON-m1, n1, o1 and p1) have a higher fraction of input from non-MB neurons than single-compartment MBONs. The fraction of inputs from PNs to MBONs via KCs is shown within the fraction of KC input (different shades of blue), computed by the product of the PN-to-KC and KC-to-MBON connections. Most MBONs receive a high fraction of olfactory input via KCs while few MBONs (b3, o1) get nearly half of their inputs via KCs from non-olfactory PNs. **c** Percentages of different types of PN input to the KCs. While there are almost equal numbers of olfactory and non-olfactory PNs synapsing onto KCs, non-olfactory PNs represent only about 14% of the inputs to KC dendrites.

We analyzed the detailed structure of KC → MBON connectivity to determine the nature of the sensory signals relayed to MBONs. About 23% of the KC input to MBONs originates in KCs that integrate inputs from non-olfactory PNs (Fig. 4b, c), significantly more than in networks with shuffled PN modalities (p < 0.05). This pattern of non-olfactory input is stereotyped: the fraction of non-olfactory input received by homologous MBONs across hemispheres is more similar than across different MBONs in the same hemisphere (p < 0.001, Mann-Whitney U Test). These observations, along with the sparse activity of olfactory KCs seen in adult flies^31^, suggest that non-olfactory signals may have a large influence on the activity of certain MBONs despite the small number of non-olfactory KCs. To quantify this influence, we compared the total number of synapses made by thermosensory KCs onto the different MBONs to 0.05 times the total number of synapses made by non-thermosensory KCs (Ext. Data Fig. 9f). This ratio quantifies the relative influence of a stimulus that activates thermosensory KCs to a typical odor stimulus that activates 5% of olfactory KCs^31^. The ratio is high for some MBONs (d3, o1 and b3) and stereotyped for homologous MBONs across hemispheres, suggesting that these MBONs may be wired to respond strongly to non-olfactory cues, such as temperature.

We also examined whether the olfactory input received by homologous MBONs is stereotyped by computing the correlation between the number of connections they receive from each olfactory PN via KCs. Unlike for non-olfactory input, homologous MBONs across hemispheres receive a less similar pattern of olfactory PN input than different MBONs in the same hemisphere (p > 0.99, Mann-Whitney U Test), arguing against stereotypy. Therefore the lack of stereotypy in the olfactory PN → KC connectivity and the greater degree of stereotypy in the non-olfactory PN → KC connectivity are inherited by the MBONs.

Interestingly, MBONs do not receive their inputs exclusively from KCs, DANs and other MBONs. We found that some compartments (Fig. 4b) have two kinds of MBONs that differ in the amount of input they receive from non-MB sources. Some MBONs are postsynaptic almost exclusively to MB neurons (over 90%), while some receive 50% or more of their inputs from non-MB neurons on dendritic branches outside the MB compartments (Fig. 4b and Supp. Table 2a). MBONs with significant input from outside the MB typically receive input from other MBONs as well. The convergence of modulatory neurons, of olfactory, thermal and visual KCs, and of other non-MB inputs onto some MBONs makes them flexible sites for learning and integration of multisensory and internal state information (via DANs, as suggested by functional studies in the adult^19;28;44;45^, and possibly via the “other” new non-MB inputs identified here).

## The MBON output network

In adult *Drosophila*, MBONs form a multi-layered feedforward network^18;28^. Consistent with this, we found synaptic connections between MBONs in different compartments, on both the ipsi- and contralateral sides, that create a bilaterally symmetric structured feedforward circuit (Fig. 4a, Supp. Table 5). Most inter-lobe connections are mediated by GABAergic (MBON-g1, g2, h1, h2) and glutamatergic (MBON-i1, j1, k1) MBONs, potentially providing a substrate for lateral inhibition between the lobes. Glutamate has previously been found to be inhibitory in the fly^46^, but an excitatory effect on some neurons cannot be excluded. Some vertical lobe (VL) MBONs could also disinhibit (via inhibition of inhibitory medial lobe (ML) MBONs), or directly excite other VL MBONs, potentially providing within-region facilitation.

In addition, there is a hierarchy of interactions across regions of the MB. MBONs of the peduncle and calyx are exclusively at the bottom of the inhibitory hierarchy, receiving inputs from GABAergic MBONs from both the VL and the ML, but not synapsing onto any other MBONs (Fig. 4a). Furthermore, the ML may also disinhibit the peduncle MBON-c1 that the VL inhibits (Fig. 4a).

## Homo- and hetero-compartment MBON connections to DANs and OANs

The mushroom body connectome further revealed several feedback connections from MBONs onto MBINs of the same compartment (Fig. 5a–b). MBON-e2 from the tip compartment of the VL (UVL) synapses onto the dendrites of OAN-e1 outside the MB (bilateral; Fig. 5a and Supp. Fig. 1b). MBON-q1 from IVL/LVL compartments is presynaptic to the axons of the DANs from these two compartments in addition to the MBIN-l1 from the LA compartment (Fig. 5b and Supp. Fig. 1b). An MBON → DAN feedback connection was found in the *α*1 compartment of the adult VL and is implicated in the formation of long-term memory^23^.

**Figure 5.**
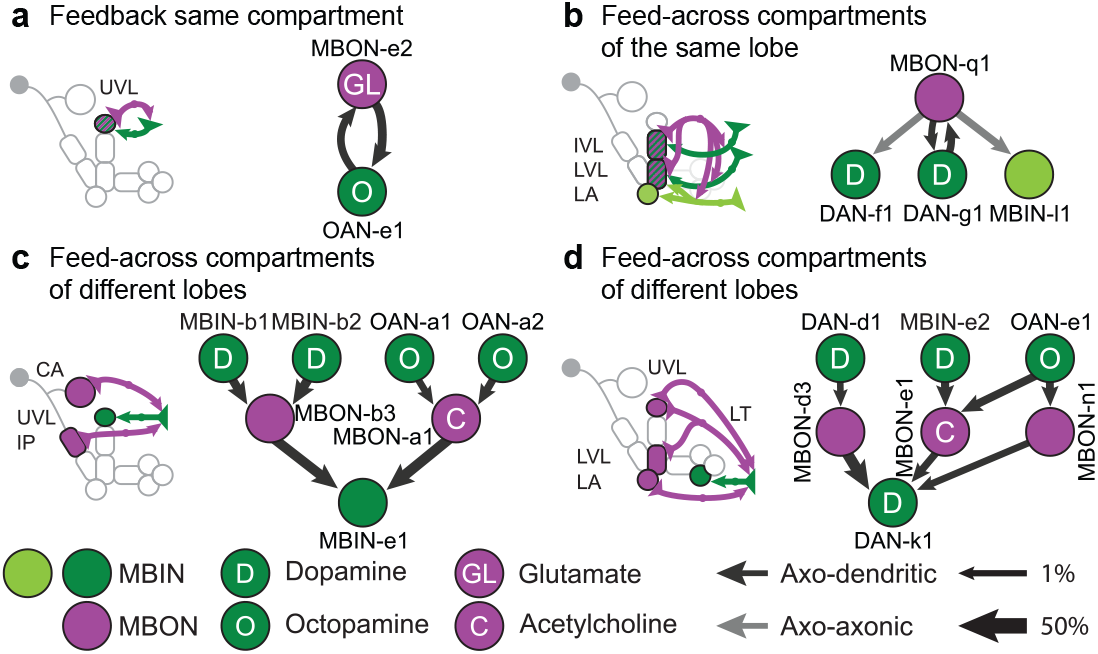
Intra- and inter-compartment feedback: MBONs of one MB compartment synapse onto MBINs of the same or other compartments. Schematics of connections from output neurons (MBONs) onto input neurons (MBINs) within their own (feedback) or from other compartments (feed-across). **a** The output neuron MBON-e2 synapses onto the dendrites of OAN-e1 in its own compartment, UVL. **b** Feedback among compartments of the same lobe suggests that the establishment of a memory in a compartment can affect the DANs of adjacent compartments. **c** Feed-across motif from proximal MB compartments (calyx and intermediate peduncle) to a distal one (UVL). **d** Feed-across motif from the vertical lobe to the medial lobe.

Interestingly, we also found hetero-compartment feed-across connections where MBONs that innervate one region of the MB synapse onto MBINs innervating other regions (Fig. 5c and Supp. Fig. 1b). The feed-across motifs could play a role during conflicting memory formation^20;41^ or during reversal learning and more generally they could enhance the flexibility of modulatory input to the MB.

## Discussion

We provide the first complete wiring diagram of a parallel fiber circuit for adaptive behavioral control. Such circuits exist in various forms, for example the cerebellum in vertebrates and the mushroom body in insects. They contribute to multiple aspects of behavioral control including stimulus classification, the formation and retrieval of Pavlovian associations, and memory-based action selection^1–6^. A comprehensive wiring diagram of such a multi-purpose structure is an essential starting point for functionally testing the observed structural connections and for elucidating the circuit implementation of these fundamental brain functions.

Even though individual neurons may change through metamorphosis, many of the basic aspects of the MB architecture are shared between larval and adult *Drosophila* stages and with other insects (see Supp. Table 6, Supp. Table 7)^1;3–6;8;18;39;40^. We therefore expect that the circuit motifs identified here are not unique to the L1 developmental stage, but instead represent a general feature of *Drosophila* and insect MBs.

## A canonical circuit in each MB compartment with unexpected motifs

Our EM reconstruction revealed a canonical circuit in each MB compartment that features two unexpected motifs in addition to the previously known MBIN → KC and KC → MBON connections. First, we were surprised to observe that the number of KC → MBIN and KC → MBON synapses are comparable. As KCs were shown to be cholinergic in adults^36^, KC → MBIN connections could be potentially depolarizing. Untrained, novel odors can activate DANs in adult *Drosophila*^47;48^ and OANs in bees^3^. Similar brief short-latency activations of dopamine neurons by novel stimuli are observed in monkeys, too, and are interpreted as salience signals^49^. Learning could potentially modulate the strength of the KC → MBIN connection, either weakening it, or strengthening it. The latter scenario could explain the increase in DAN activation by reinforcement-predicting odors observed in adult *Drosophila* ^47^, bees and monkeys^3;49^. In addition, dopamine receptors have been shown to be required in *Drosophila* KCs for memory formation^4^. Another unexpected finding was that MBINs synapse directly onto MBONs, rather than only onto KCs. Such a motif could provide a substrate for neuromodulator-gated Hebbian spike-timing dependent plasticity which has been observed in locust MB^38^.

## Single-claw KCs and the dimensionality of the MB representation

In addition to random and bilaterally asymmetric olfactory and structured non-olfactory PN → KC connectivity (Ext. Data Fig. 3), our analysis identified single-claw KCs whose number and lack of redundancy are inconsistent with random wiring (Fig. 2). Random wiring has previously been shown to increase the dimension of sensory representations when the number of neurons participating in the representation is large compared to the number of afferent fibers, as in the cerebellum or adult mushroom body^13;33^. However, our model shows that when the number of neurons is limited, random wiring alone is inferior to a combination of random and structured connectivity that ensures each input is sampled without redundancy. The presence of single-claw KCs may reflect an implementation of such a strategy. In general, our results are consistent with a developmental program that produces complete and high-dimensional KC sensory representations to support stimulus discrimination at both larval and adult stages.

## Mutual inhibition as general motif of the MBON-MBON network

This study reveals the complete MBON-MBON network at synaptic resolution (Fig. 4a). Previous studies in the larva have shown that odor paired with activation of medial and vertical lobe DANs leads to learned approach^8^ and avoidance^7;42^, respectively. Our connectivity analysis reveals that glutamatergic MBONs from the medial lobe laterally connect to MBONs of the vertical lobe. The glutamatergic MBON → MBON connections could be inhibitory^46^, although further studies are needed to confirm this. Furthermore, inhibitory GABAergic MBONs from the vertical lobe laterally connect to MBONs of the medial lobe. An example is the feedforward inhibition of ML MBON-i1 output neuron by the VL GABAergic MBON-g1, -g2 output neurons. A similar motif has been observed in *Drosophila* adult, where aversive learning induces depression of conditioned odor responses in the approach-promoting MBON-MVP2 (MBON-11), which in turn disinhibits conditioned odor responses in the avoidance-promoting MBON-M4/M6 (MBON-03) due to the MBON-MVP2 to MBON-M4/M6 feedforward inhibitory connection^18;25–28;40^.

Combining the present connectivity analysis of the MBON-MBON network in the larva and previous studies in the adult *Drosophila*^5;18;25;28;40^, the rule seems to be that MBONs encoding opposite learnt valance laterally inhibit each other. Such inhibitory interactions have been proposed as a paradigmatic circuit motif for memory-based action selection^50^.

## Acknowledgements

We thank A. Khandelwal, J. Lovick, J. Valdes-Aleman, I. Larderet, V. Hartenstein, A. Fushiki, B. Afonso, P. Schlegel and M. Berck for reconstructing 31% of arbor cable and 19% of synapses. We thank R. Axel, G. M. Rubin and Y. Aso for their comments on the manuscript. ALK was supported by NIH grant #F32DC014387. AL-K and LFA were supported by the Simons Collaboration on the Global Brain. LFA was also supported by the Gatsby, Mathers and Kavli Foundations. CEP and YP were supported by the DARPA XDATA program (AFRL contract FA8750-12-2-0303) and the NSF BRAIN EAGER award DBI-1451081. KE and AST thank the Deutsche Forschungsgemeinschaft, TH1584/1-1, TH1584/3-1; the Swiss National Science Foundation, 31003A_132812/1; the Baden Württemberg Stiftung; Zukunftskolleg of the University of Konstanz and DAAD. BG and TS thank the Deutsche Forschungsgemeinschaft, CRC 779, GE 1091/4-1; the European Commission, FP7-ICT MINIMAL. We thank the Fly EM Project Team at HHMI Janelia for the gift of the EM volume, the Janelia Visiting Scientist program, the HHMI visa office, and HHMI Janelia for funding.

## Author contributions

K.E., F.L., A.L.-K., B.G., L.F.A., A.T., M.Z. and A.C. conceived the project, analyzed the data and wrote the manuscript. K.E., F.L., I.A., C.S.-M., T.S. A.T. and A.C. reconstructed neurons. K.E. performed learning experiments. A.L.-K. built the models. J.W.T contributed GAL4 lines and their imagery. R.D.F. generated EM image data. A.L.-K., C.S.-M., Y.P. and C.E.P. analyzed connectivity patterns. F.L. and A.H. performed immunostainings. C.E. generated functional data.

## Author information

The authors declare no competing financial interests. Correspondence and requests for materials should be addressed to L.F.A, A.T., M.Z. or A.C.

## METHODS

### Circuit mapping and electron microscopy

We reconstructed neurons and annotated synapses in a single, complete central nervous system from a 6-h-old [*iso*] *Canton S G1 x w*^1118^ *[iso] 5905* larva acquired with serial section transmission EM at a resolution of 3.8 x 3.8 x 50 nm, that was first published in^11^ along with the detailed sample preparation protocol. Briefly, the CNS of 6-h-old female larvae were dissected and fixed in 2% gluteraldehyde 0.1 M sodium cacodylate buffer (pH 7.4) to which an equal volume of 2% OsO_4_ in the same buffer was added, and microwaved at 350-W, 375-W and 400-W pulses for 30 sec each, separated by 60-sec pauses, and followed by another round of microwaving but with 1% OsO_4_ solution in the same buffer. Then samples were stained en bloc with 1% uranyl acetate in water by microwave at 350 W for 3x3 30 sec with 60-sec pauses. Samples were dehydrated in an ethanol series, then transferred to propylene oxide and infiltrated and embedded with EPON resin. After sectioning the volume with a Leica UC6 ultramicrotome, sections were imaged semi-automatically with Leginon^51^ driving an FEI T20 TEM (Hillsboro), and then assembled with TrakEM2^52^ using the elastic method^53^.

To map the wiring diagram we used the web-based software CATMAID^54^, updated with a novel suite of neuron skeletonization and analysis tools^55^, and applied the iterative reconstruction method^55^. All annotated synapses in this wiring diagram fulfill the four following criteria of mature synapses^11;55^: (1) There is a clearly visible T-bar or ribbon. (2) There are multiple vesicles immediately adjacent to the T-bar or ribbon. (3) There is a cleft between the presynaptic and the postsynaptic neurites, visible as a dark-light-dark parallel line. (4) There are postsynaptic densities, visible as black streaks hanging from the postsynaptic membrane.

In this study we validate the reconstructions as previously described^11;55^, a method successfully employed in multiple studies^11;55–60^. Briefly, in *Drosophila*, as in other insects, the gross morphology of many neurons is stereotyped and individual neurons are uniquely identifiable based on morphology^61–63^. Furthermore, the nervous system in insects is largely bilaterally symmetric and homologous neurons are reproducibly found on the left and the right side of the animal. We therefore validated MBON, DAN, OAN, MBIN, APL and PN neuron reconstructions by independently reconstructing synaptic partners of homologous neurons on the left and right side of the nervous system. By randomly re-reviewing annotated synapses and terminal arbors in our dataset we estimated the false positive rate of synaptic contact detection to be 0.0167 (1 error per 60 synaptic contacts). Assuming the false positives are uncorrelated, for an n-synapse connection the probability that all n are wrong (and thus that the entire connection is a false positive) occurs at a rate of 0.0167^*n*^. Thus, the probability that a connection is false positive reduces dramatically with the number of synaptic connections contributing to that connection. Even for n = 2 synapse connections, the probability that the connection is not true is 0.00028 (once in every 3,586 two-synapse connections) and we call connections with two or more connections ‘reliable’ connections. See^11;55^ for more details.

### Identification of PNs

The olfactory PNs from this same EM volume were previously traced and identified^12^, with each olfactory glomerulus being innervated by exactly one single uniglomerular PN in the larva^64^. To identify the photosensory and thermosensory PNs connected to KCs, we identified all neurons downstream of 12 photosensory neurons ^65^ in the left and right brain hemispheres and all neurons downstream of three previously characterized cold sensing neurons, expressed in 11F02-GAL4 line^66^.

### Learning in first instar larvae with a substitution experiment

Learning experiments were performed as described in^8;42;67^. The fly strain *PGMR58E02-GAL4attP2* (Bloomington Stock Center no. 41347) and the *attP2* control strain were crossed to *P20XUAS-IVS-CsChrimson.mVenusattP18* (Bloomington Stock Center no. 55134). Flies were reared at 25°C in darkness on 4% agarose with a yeast and water paste including retinal in a 0.5 mM final concentration. First instar feeding-stage larvae in groups of 30 individuals were placed on plates filled with 4% agarose and the odor ethyl-acetate (100-times diluted in distilled water) was presented on filter papers located on the lid. Larvae were exposed to constant red light (630nm, power: 350μW/cm2) during this odor presentation for 3 minutes. Subsequently larvae were transferred to a new plate and no odor was presented in the dark for 3 minutes. This paired training cycle was repeated two times. We also performed the unpaired experiment (odor presentation in the dark and red light without odor) with another group of 30 naïve larvae. After a 5-minute test with odor presentation on one side of the lid larvae were counted on the side of the odor, no odor and a 1 cm area in the middle of the plate. Preference and performance indices were calculated as in^8^. Namely for both the paired and unpaired, the number of larvae on the no-odor side are subtracted from the number of larvae on the odor side, and the result is divided by the total number of larvae on the plate (including those in the middle). The performance index is half of the value for the paired minus the unpaired preference scores.

### Random models of PN → KC connectivity

To generate random PN → KC models, individual PN connection probabilities were computed as the number of multi-claw KCs contacted divided by the total number of connections onto multi-claw KCs. Probabilities were computed separately for the two hemispheres. To generate the random networks in Fig. 2a, we assumed that PN → KC connections were formed independently according to these probabilities. We then iteratively generated KCs until the number of multi-claw KCs matched the number found in the reconstruction. To compare dimension and classification performance for random networks with varying degrees of connectivity (Fig. 2c), we scaled the individual PN connection probabilities so that, on average, each KC received between 1 and 10 claws (Fig. 2b, c). In this case, we fixed the total number of KCs (single- and multi-claw) to be equal to the number found in the reconstruction.

To obtain KCs with a specified number of claws (Fig. 2b), PN connections were determined by weighted random sampling using the individual connection probabilities described above. When comparing detailed connectivity statistics to random models (Ext. Data Fig. 3), we used this type of sampling to match the KC claw count distribution for the random models to that of the data.

We also evaluated how many random multi-claw KCs would be required to ensure coverage of the subset of PNs that are connected to single-claw KCs. We restricted our analysis to these PNs and iteratively added random multi-claw KCs, with the claw counts distribution determined by the reconstructed connectivity, until each PN was connected to at least one KC. For this analysis, PN connection probabilities were computed using the full PN → KC connectivity matrix.

### Model of sparse KC responses

We assumed that the activity of the ith KC was given by *s_i_* = [Σ_*j*_ *J_ij_x_j_* – *θ_i_*]+, where [·]_+_ denotes rectification, *J* is the matrix of PN → KC connections, *x_j_* is the activity of the *j*th PN, and *θ_i_* is the KC activity threshold. Simulated odor-evoked PN activity was generated by sampling the value of each *x_j_* independently from a rectified unit Gaussian distribution. The value of *θ_i_* was adjusted so that each KC fired for a fraction *f* = 0.05 of odors. The entries of *J* were either the synapse counts from the reconstructed data or, for the case of random connectivity, chosen in the manner of the previous section with a weight sampled randomly from the distribution of PN → multi-claw-KC synaptic contact numbers found in the reconstruction.

To assess classification performance (Fig. 2b, c), the KC responses to 8 odors were evaluated (10 odors were used when simulating the adult MB). Odor responses again consisted of rectified unit Gaussian random variables, but corrupted by Gaussian noise with standard deviation 0.2. A maximum-margin classifier was trained to separate the odors into two categories to which they were randomly assigned, and the error rate was assessed when classifying odors with different noise realizations. Dimension (Fig. 2f) was assessed by estimating the KC covariance matrix *C_ij_* = 〈(*z_i_* – 〈*z_i_*〉)(*z_j_* – 〈*z_j_*〉)〉, where *z_i_* is the z-scored activity of neuron *i*, using 1,000 random simulated odors. We then computed 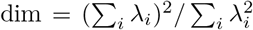, where {λ_*i*_} are the eigenvalues of *C*^33^.

To implement recurrence, we modified the model so that the activity of the *i*th KC was given by 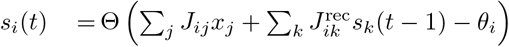, where 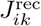 is a matrix of recurrent KC → KC interactions and *t* is the timestep. *J*^rec^ is equal to *α*_1_*J*^KG→KG^ + *α*_2_*J*^APL^, where *J*^KG→KG^ was determined by the KC → KC synapse counts (either dendro-dendritic only or both dendro-dendritic and axo-axonic) and 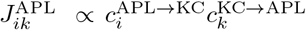, with 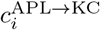 representing the number of APL synapses onto the *i*th KC and 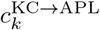 representing the number of synapses onto APL from the *k*th KC. *J*^APL^ was scaled so that the average of its entries equaled that of *J*. The scalar *α*_1_ = 0.33 was chosen so that the strength of KC → KC and PN → KC connections were comparable, while *α*_2_ = 0.05 represents the gain of the APL neuron. A single randomly chosen KC was updated on each timestep, with the number of timesteps equal to 5 times the number of KCs. We also modeled facilitatory KC → KC interactions by assuming KC → KC connections only depolarize the postsynaptic neuron when it is already above threshold.

### Immunostaining

Third-instar larvae were put on ice and dissected in PBS. For all antibodies brains were fixed in 3.6% formaldehyde (Merck) in PBS for 30 min except anti-dVGlut that required bouin’s fixation^68^. After several rinses in PBT (PBS with 1% or 3% Triton X-100; Sigma-Aldrich), brains were blocked with 5% normal goat serum (Vector Laboratories) in PBT and incubated for at least 24 hours with primary antibodies at 4°C. Before application of the secondary antibodies for at least 24 hours at 4°C or for 2 hours at room temperature, brains were washed several times with PBT. After that, brains were again washed with PBT, mounted in Vectashield (Vector Laboratories) and stored at 4°C in darkness. Images were taken with a Zeiss LSM 710M confocal microscope. The resulting image stacks were projected and analyzed with the image processing software Fiji^69^. Contrast and brightness adjustment, rotation, and arrangement of images were performed in Photoshop (Adobe Systems).

### Antibodies

Brains were stained with the following primary antibodies, polyclonal goat anti-GFP fused with FITC (1:1000, Abcam, ab6662), polyclonal rabbit anti-GFP (1:1000, Molecular Probes, A6455), polyclonal chicken anti-GFP (1:1000, Abcam, ab13970), monoclonal mouse anti-TH (1:500, ImmunoStar, 22941), polyclonal rabbit anti-TDC2 (1:200, CovalAb, pab0822-P), monoclonal mouse anti-ChAT (1:150, Developmental Studies Hybridoma Bank, ChaT4B1), polyclonal rabbit anti-GABA (1:500, Sigma, A2052), and rabbit anti-dVGlut (1:5000)^70^ for identifying GFP positive, dopaminergic, octopaminergic, cholinergic, GABAergic, and glutamatergic neurons, respectively. The following secondary antibodies were used, polyclonal goat antichicken Alexa Fluor 488 (1:200, Molecular Probes, A11039), polyclonal goat anti-rabbit Alexa Fluor 488 (1:200, Molecular Probes, A11008), polyclonal goat anti-rabbit Alexa Fluor 568 (1:200, Molecular Probes, A11011), polyclonal goat anti-rabbit Cy5 (1:200, Molecular Probes, A10523), polyclonal goat antimouse Alexa Fluor 647 (1:200, Molecular Probes, A21235), and polyclonal goat anti-rabbit Alexa Fluor 647 (1:200, Molecular Probes, A21245).

### Identifying GAL4 lines that drive expression in MB-related neurons

To identify GAL4 lines (listed in Supp. Table 3) that drive expression in specific MB-related neurons, we performed single-cell FLP-out experiments (for flp-out methodology see^11;71^) of many candidate GAL4 lines (LiTruman2014). We generated highresolution confocal image stacks of the projection patterns of individual MBONs/DANs/OANs (multiple examples per cell type), which allowed their identification. Most MBONs/DANs/OANs were uniquely identifiable based on the dendritic and axonal projection patterns (which MB compartment they project to and the shape of input or output arbour outside the MB). These were also compared to previously reported single-cell FLP-outs of dopaminergic and octopaminergic neurons in the larva^8;9;72–74^. Some compartments were innervated by an indistinguishable pair of MBONs or MBINs.

### Estimating the birth order of KCs

KCs arise from four neuroblasts per hemisphere that divide continuously from the embryonic to the late pupal stage^75;76^. The primary neurite of newborn KCs grows through the center of the MB peduncle, pushing existing KCs to the surface of the peduncle^77^ (Ext. Data Fig. 4h). As new cells are added, their somas remain closer to the neuroblast at the surface of the brain and push away existing somas. The point of entry into the lineage bundle of each KC primary neurite remains as a morphological record of the temporal birth order, with some noise. For every KC, we measured the distance from the point where its primary neurite joins the lineage bundle to a reference point consisting of the point where the neurite of the last KC (potentially the oldest of that lineage) joins the bundle before the bundle enters the neuropil (Fig. 1f) to form the peduncle together with the other three bundles. This measurement revealed that single-claw KCs are born early in MB development, followed by 2-claw KCs, 3-claw KCs and so on (Fig. 1f), with the last-born KCs being immature and expected to develop into multi-claw KCs later in larval life^78^.

### Data availability

The EM volume is available at http://openconnecto.me/catmaid/, titled “acardona_0111 _8”. The coordinates for the skeletons modeling neuronal arbors and for the synapses, and the data tables from which all graphs were made, are in the supplementary material.

